# Not all seed transfer zones are created equal: Using fire history to identify seed needs in the Cold Deserts of the Western US

**DOI:** 10.1101/2022.08.15.503985

**Authors:** Sarah C. Barga, Francis F Kilkenny, Scott Jensen, Sarah M. Kulpa, Alison C. Agneray, Elizabeth A. Leger

## Abstract

Restoration planning requires a reliable seed supply, yet many projects occur in response to unplanned events. Identifying regions of greater risk could help guide seed procurement. Using fire perimeters (2000–2019), we investigated differences in fire occurrence (frequency, area burned, percent of area burned) among seed transfer zones within Cold Deserts of the US. We considered both provisional seed transfer zones (PSTZs), created using climate and stratified by ecoregion, and empirical seed transfer zones (ESTZs) for two species commonly used in restoration. Finally, we present a case study on fire occurrence within Northern Basin and Range greater sage-grouse Priority Areas for Conservation (PACs). Historic fire was effective for prioritizing seed zones: 23 of 132 PSTZs burned every year, and, within each ecoregion, two PSTZs comprised ≧ 50% of the total area burned across all years. Similarly, fire disproportionately occurred in some ESTZs; in the Northern Basin and Range, one (*Artemisia tridentata* - 83%) or two zones (*Pseudoroegneria spicata* - 65%) made up a majority of total area burned. Fire occurrence within PACs largely reflected the PSTZ and ESTZ priorities found for the ecoregion, with small exceptions. Imperiled PSTZs (high proportion burned) in PACs largely reflected the patterns found across the ecoregion, while imperiled ESTZs departed from ecoregional patterns. Considering historic disturbance can focus seed procurement efforts on regions that encounter regular disturbance, experience large disturbances, or have particular conservation value. This information can guide seed production, purchase, and storage, create more certainty for growers and managers, and ultimately increase restoration success.

**IMPLICATIONS FOR PRACTICE:**

- Available data on past disturbance patterns may inform strategies for prioritizing seed procurement decisions, especially as geospatial information becomes more widely available
- The methods presented here illustrate an approach for using historic disturbance data to identify regions that are of greatest need for seed collection and conservation, using seed transfer zones within US Cold Desert ecoregions as an example
- Identifying regions that experience disturbance over large areas or are likely to be imperiled due to disturbance at a high proportion can guide the collection and conservation of plant materials and better align available plant material with future restoration needs

## INTRODUCTION

Predicting seed needs for restoration is a worldwide challenge (Breed et al. 2018), and access to desired seeds is a common limitation for restoration projects (Broadhurst et al. 2008; Harrison et al. 2020). Whether wild-collected or sourced from agricultural production fields, the complexity of plant life histories requires advanced planning to ensure that seeds are available when they are needed (Erickson & Halford 2020; Pedrini et al. 2020). When restoration projects occur in response to planned disturbances, proactive management can ensure that seeds are stored and available when needed, by conserving topsoil and associated seed banks (Golos et al. 2016) or collecting and producing seeds in anticipation of use (Erickson & Halford 2020). However, many restoration projects are conducted in response to disturbance events, such as hurricanes, floods, or fires (BLM 2007). The uncertain timing, size, and location of this type of restoration makes ensuring seed availability a challenge for such projects. By using historic information and prediction of future events to forecast needs, it may be possible to proactively ensure that required seed resources are available for the areas most likely to need them (Harrison et al. 2020).

Wildland fire is an unplanned disturbance that motivates restoration projects in many regions around the world. With climate change and other anthropogenic disturbances, even fire-adapted ecosystems may require active restoration to return to desirable states (Fulé 2008). For many plant communities, the question is not whether they will burn, but when (Pitman et al. 2007; Liu & Wimberly 2016), and what type of management may be needed to maintain desired successional trajectories (Ott et al. 2022). Sagebrush steppe shrubland, the predominant vegetation type in the western United States Cold Deserts, is made up of plant communities that are vulnerable and maladapted to increased wildfire (Winward 1985). Over 40% of the sagebrush steppe Cold Desert has significant cover of invasive annual grasses (Larson and Tuor 2021), such as cheatgrass (*Bromus tectorum*), which drive fire cycles and outcompete native vegetation under higher disturbance regimes (D’Antonio & Vitousek 1992; Pilliod, Welty & Arkle 2017). Depending on site conditions, wildfire in invaded areas is between 2.5 and 22 times more frequent than it was prior to European settlement (Whisenant 1990; Balch et al. 2013). Once covering more than 600,000 km^2^, 60–90% of the sagebrush steppe has been lost, fragmented, or degraded (Noss et al. 1995; Knick et al. 2003). Due to these disturbances, the sagebrush steppe ecosystem is facing a loss in native plant diversity (Mahood & Balch 2019), which results in a loss of evolutionary history and potential of native plants (McArthur & Fairbanks 2001), along with reduced ecosystem services such as water and soil retention and productivity (Kachergis et al. 2011; Nichols et al. 2021). These changes also threaten animal populations, including species of conservation concern, such as the greater sage-grouse (*Centrocercus urophasianus*) (Knick et al. 2003).

To counteract the effects of wildfire and exotic plant invasions, land management agencies, such as the US Department of Interior, Bureau of Land Management (BLM), commonly seed after fire in the sagebrush steppe. The BLM has seeded after fire since the 1940s, with significant increases in yearly seeding occurring in the 1980s and 2000s in response to increasing fire frequencies (Pilliod, Welty & Toevs 2017). Between 2015 and 2020, the BLM spent $222M on emergency stabilization and burned area rehabilitation, including $67M on seed procurement, most of which was used in the sagebrush steppe (BLM 2016–2021). Nevertheless, short-term targets for soil stabilization and annual grass suppression have met with mixed success (Knutson et al. 2014), and attempts at establishing diverse plant communities that support wildlife populations of conservation concern have generally been unsuccessful (Arkle et al. 2014). In addition to the challenges of restoring heavily invaded arid sites, these outcomes are likely due to the prominent use of non-native Eurasian bunchgrasses, which establish well and can outcompete annual grasses (Ott et al. 2019), but interfere with native species recovery (Nafus et al. 2015; Williams et al. 2017; Ott et al. 2022).

However, even when native species are used, which is occurring with increasing frequency (Pilliod, Welty & Toevs 2017), native seed sources may fail to establish and thrive due in part to a mismatch between the source environment, where the population has been subject to natural selection, and the environment at the restoration site (McKay et al. 2005). This may be especially true in sagebrush steppe ecosystems, where plant populations are often adapted to local environments (Baughman et al. 2019).

Several nations have policies and strategies that call for the use of seed transfer zones (STZs) to guide the use of native plant seed sources for use in restoration (e.g., Prasse et al. 2010). The National Seed Strategy for Rehabilitation and Restoration (PCA 2015) provides this guidance in the United States, and state and regional frameworks have been developed for the sagebrush steppe ecosystem (e.g., NNSP 2020). STZs are designed to limit the chances that seed sources are maladapted to restoration site conditions and can help identify populations important for genetic conservation (Kilkenny 2015; Massatti et al. 2018). Ideally, species-specific empirical STZs (ESTZs) would be used for each species within a seed mix (Johnson et al. 2010). ESTZs are being developed for “workhorse” restoration species in the sagebrush steppe, such as bluebunch wheatgrass (*Pseudoroegneria spicata*) (St. Clair et al. 2013) and sagebrush (Richardson & Chaney 2018). However, given the number of species that are important to ecosystem resiliency in the sagebrush steppe and other ecosystems (Dumroese et al. 2015), developing ESTZs for every species of interest is impractical. Generalized provisional STZs (PSTZs) have been developed for use in restoration seed sourcing in the United States when ESTZs are not available (Bower et al. 2014). These PSTZs are based on two climate variables, winter minimum temperature and annual heat moisture index, that are adaptively important across plant taxa, and can be stratified by Omernik’s level III ecoregions to further minimize instances of maladaptation (Omernik 1987; Bower et al. 2014).

The topographic and climatic complexity of the sagebrush steppe generates a relatively high number of PSTZs within relatively small geographic areas. This creates a helpful framework for seed need planning, and can create opportunities to make the seed procurement process more efficient for land managers, seed collectors, and seed producers (Taylor et al. 2018). As a Cold Desert, the sagebrush steppe has low productivity and, while wild collection is possible for some species (e.g., widespread and abundant shrubs like *A. tridentata*), for many species, wildland seed cannot be collected in high enough quantities to fulfill restoration needs. Therefore, seed must be increased from initial collections using agricultural production methods (Pedrini et al. 2020). This process takes planning, funding, and logistic coordination over multiple years and reduces flexibility in matching genetically appropriate seed to unplanned disturbance events and restoration projects. The majority of plant materials used in sagebrush steppe restoration were developed through rangeland improvement programs, some beginning in the 1930s, that maximized forage production (Vallentine 1989) over genetic appropriateness or ecological compatibility with local plant communities (Leger & Baughman 2015). Given the practical constraints of developing and producing genetically appropriate plant materials for post-fire restoration in the sagebrush steppe, ways to prioritize these processes are needed.

Considering fire history and risk in the sagebrush steppe ecosystem could be used to prioritize seed needs. While fire forecasting models are improving (e.g., Krawchuk et al. 2009), we have now amassed sufficient fire history for this region to reveal patterns that can potentially be used as an indication of likely seed needs for the future (Fig. 1). This approach, where past disturbance is used to forecast future need, could be applied across PSTZs, for all species, as well as across ESTZs for workhorse restoration species. Knowledge of likely seed needs would allow for the prioritization of seed collections for long-term genetic conservation and seed production for use in post-fire restoration. This could create reliable demand for seed growers, increase availability of desirable seed, and lead to improved post-fire restoration outcomes in the western US.

**Figure 1.**
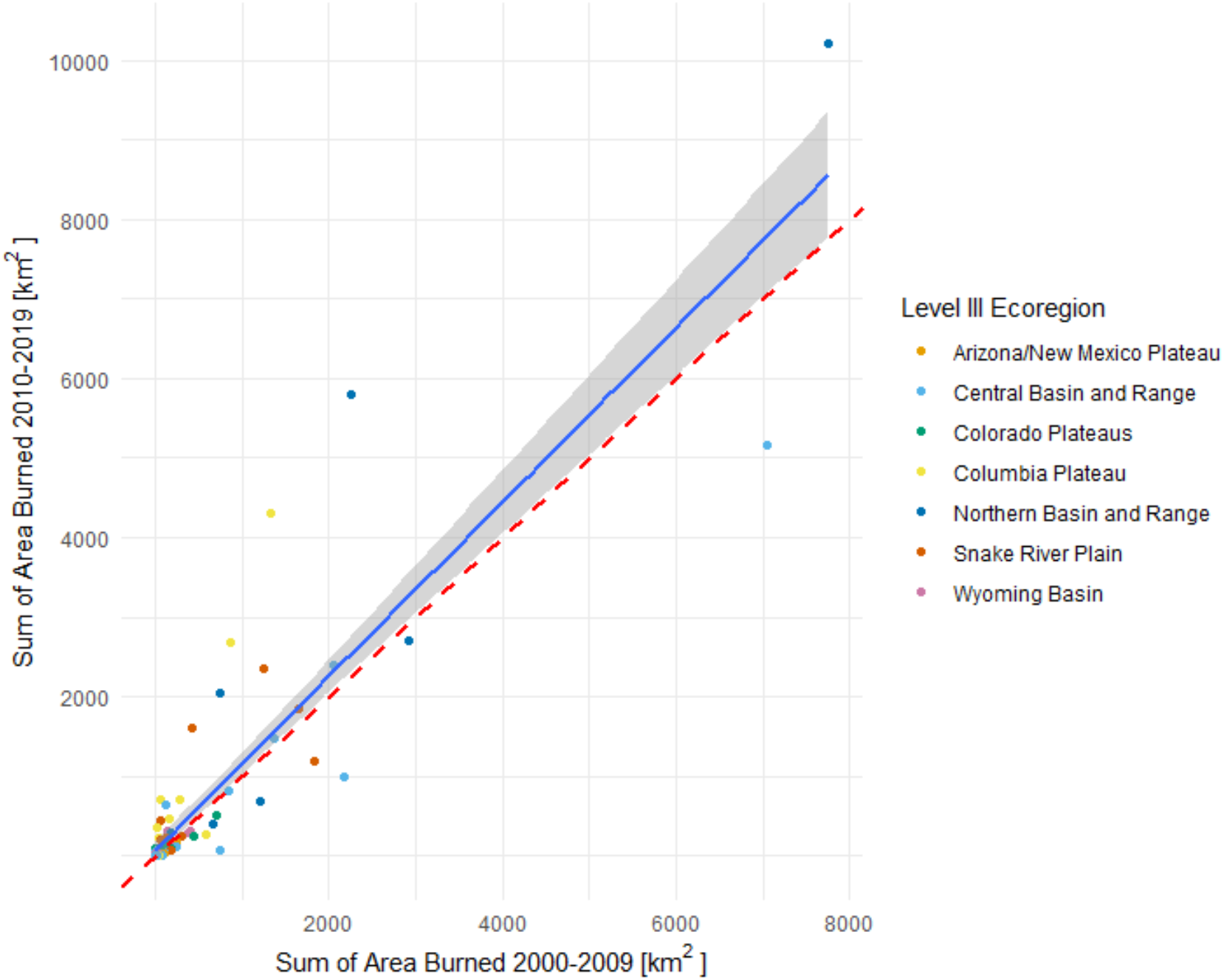
Comparison of the sum of the area burned for each provisional seed transfer zone (PSTZ) within each Omernik Level III ecoregion from 2000–2009 with the sum of the area burned from 2010–2019. The red dashed line represents the trendline if there were equal levels of burning within the two time periods, with points above the dashed line representing areas experiencing more burning than in the past. The actual trendline, in blue, represents the pattern of burning across all PSTZs, with 95% confidence interval in gray, showing that past levels of burning are generally predictive of future levels of burning.

Here, we consider fire history between 2000–2019 in the western US Cold Deserts, presenting an example of how seed needs can be prioritized in an imperiled landscape. We use three components to assess the occurrence of fire in STZs within Cold Deserts and wildlife management areas: the frequency of fire (measured as the number of years an area has burned in our 20-year timeframe), the total area burned, and the percent of area burned (to identify potentially imperiled STZs). We first consider fire occurrence within PSTZs, answering the following specific question:

1.What are the patterns of wildfire occurrence within Cold Desert ecoregions of the western US, with a focus on PSTZs?

Next, we present two case studies on the ways this information could be used to prioritize seed development, either A) within ecoregions or B) within wildlife habitat of particular value. For this analysis, in addition to PSTZs, we use ESTZs that have been developed for two foundational species, bluebunch wheatgrass (*Pseudoroegneria spicata;* typically agriculturally produced) and Wyoming and basin big sagebrush (*Artemisia tridentata* ssp. *wyomingensis* and *tridentata*; typically wild collected). For A), we focus on the Northern Basin and Range, an ecoregion with extensive fire, and for B), we focus on the greater sage-grouse Priority Areas for Conservation (PACs), a designation of management units identified in the 2013 greater sage-grouse Conservation Objectives Team Report (USFWS 2013), which overlaps the Northern Basin and Range and is one of the most burned regions for this species of conservation concern. In these case studies, we address the following questions:

2.What are the patterns of wildfire occurrence in the Northern Basin and Range, with a focus on ESTZs for *A. tridentata* and *P. spicata*?

3.What are the patterns of wildfire occurrence in greater sage-grouse PACs within the Northern Basin and Range ecoregion, looking at both PSTZs and ESTZs within that area?

Based on our observations of wildfire occurrence in the Great Basin, we expected that fires would be concentrated in particular PSTZs, rather than equally spread across Cold Deserts. Further, we expected that, within the Northern Basin and Range, we would be able to identify priority STZs for seed needs for our focal species, which would help guide seed development. Finally, we expected that there would be overlap between the priority STZs identified in questions 1 and 2 and the occurrence of fire within priority management areas representing important greater sage-grouse conservation areas.

Through this work, we aim to illustrate how to identify seed collection locations that are likely to be needed for restoration, encompassing multiple species and habitat management goals.

## METHODS

### Study Area

Environmental Protection Agency (EPA) defined ecoregions are areas that have similar biotic, abiotic, terrestrial, and aquatic ecosystem components and are designed to provide a meaningful spatial framework for assessing ecosystem management strategies (Omernik 1987). Our study focuses on the EPA Level II ecoregion defined as Cold Desert, which is composed of seven EPA Level III ecoregions (Table 1). These areas are generally defined as having hot, dry summers and cold or mild winters, with annual average precipitation that ranges from 4 mm in lower elevation basins to as much as 1000 mm at higher elevations (Table 1). Vegetation is made up primarily of arid sagebrush steppe, grassland, or salt desert scrub plant communities.

**Table 1.** Descriptions of Omernik Level III ecoregional characteristics within Cold Deserts *

| Ecoregion | Location/Elevation | Dominant Vegetation | Climate |
| --- | --- | --- | --- |
| Columbia Plateau | Central and southeastern Washington, north-central Oregon, and a small part of northwestern Idaho; elevation 60–1,500 m | Sagebrush steppe and grassland | Hot, dry summers; cold winters; MAT 7–12 °C, frost-free period 70–190 d; MAP 334 mm, range: 150–600 mm |
| Wyoming Basin | Central and western Wyoming with small extensions into Montana, Colorado, Utah, and Idaho; elevation 1,220–2,850 m | Arid grasslands and shrublands with some pinyon-juniper woodland | Warm-hot summers; cold winters; MAT 0–8 °C, frost-free period 30–130 d; MAP 296 mm, range: 130–500 mm |
| Northern Basin and Range | Northern Great Basin, including: southeast Oregon, northern Nevada, southern Idaho, and small portion of northern Utah; elevation 800–3,000 m | Low elevations: sagebrush steppe and cool season grasses, high elevations: Douglas fir and aspen | Hot summers; cold winters; MAT 5–9 °C, frost-free period 30–140 d; MAP 351 mm, range: 150–1000 mm |
| Snake River Plain | Southern Idaho; elevation 640–1,980 m | Sagebrush steppe | Warm summers; cold winters; MAT 6–10 °C, frost-free period 50–170 d; MAP 316 mm, range: 110–650 mm |
| Central Basin and Range | Central Great Basin, including: Nevada, western Utah, and small extensions into California and southern Idaho; elevation 1,020–4,000 m | Sagebrush and saltbush-greasewood vegetation with some pinyon-juniper and Douglas fir and aspen | Hot summers; mild winters; MAT 2–14 °C, frost-free period 15–200 d; MAP 277 mm, range: 4–1000 mm |
| Colorado Plateaus | Eastern and southern Utah, western Colorado, and small portions of northern Arizona and northwestern New Mexico; elevation 900–3,000 m | Blackbrush-saltbush, sagebrush, and pinyon-juniper | Hot, dry summers; cold, dry winters; MAT 5–15 °C, frost-free period 50–220 d; MAP 298 mm, range: 130–800 mm |
| Arizona/New Mexico Plateau | Northern Arizona, northwestern New Mexico, and southern Colorado; elevation 1,370–3,000 m | Saltbush-greasewood vegetation, sagebrush, and pinyon-juniper | Hot, dry summers; cold, dry winters; MAT 5–16 °C, frost-free period 50–250 d; MAP 293 mm, range: 125–380 mm |

\* This information was summarized from the US Level III Ecoregion Descriptions prepared by the Commission for Environmental Cooperation (CEC) and can be found at https://www.epa.gov/eco-research/ecoregions-north-america; MAT = mean annual temperature, MAP = mean annual precipitation

### Geospatial Data

All geospatial data processing was performed in ArcMap 10.5.1 (ESRI 2016). We used fire perimeters from the Geospatial Multi-Agency Coordination Wildfire Application (GeoMac) to represent fire perimeters within our analyses; GeoMac data are based on annual fire data and contain shapefiles with perimeters for individual fires recorded as separate features. Fire perimeters were downloaded for each fire year (2000–2019, Accessed: March 9, 2020), and the Dissolve function (Data Management toolbox, Generalization toolset) was used to aggregate all individual fire features into a single feature representing fires for a particular year.

Geospatial data for provisional STZs for the United States and empirical STZs for *A. tridentata* (ssp. *wyomingensis* and *tridentata)* and *P. spicata* were obtained from the Western Wildland Environmental Threat Assessment Center (WWETAC) website (https://www.fs.fed.us/wwetac/threat-map/TRMSeedZoneData.php). Bower et al. (2014) delineated discrete PSTZs for the continental US using a combination of winter minimum temperature and aridity, measured as the annual heat:moisture index (AH:M) and calculated as mean annual temperature plus 15 °C divided by mean annual precipitation in meters—larger numbers equate to higher aridity. We further stratified the original 64 PSTZs created by Bower, using Omernik Level III ecoregions, to create a total of 132 PSTZs considered in our analyses. Richardson and Chaney (2018) created ESTZs for *A. tridentata* using flowering phenology and survival from common gardens and climate from seed source locations to identify discrete STZs using a numeric naming convention. St. Clair et al. (2013) described ESTZs for *P. spicata* using size, flowering phenology, and leaf width from common gardens and climate from seed source locations to delineate discrete STZs using an alpha-numeric naming convention.

Geospatial data for Sage-Grouse Priority Areas for Conservation (PACs) were obtained by request from the US Fish and Wildlife Service and the US Geological Survey. These data are described in the 2013 greater sage-grouse Conservation Objectives Team (COT) Report (USFWS 2013). The PACs represent areas identified by individual states as essential for the long-term conservation of sage-grouse. The COT has concluded that the PACs are key for range-wide species conservation.

### Quantifying Fire Occurrence

We quantified fire occurrence in different PSTZs and ESTZs within Cold Desert Level III ecoregions and a priority sage-grouse conservation area using three components: burn frequency, the total area burned, and the percent of total area burned. We calculated burn frequency by overlaying all twenty years of fire data and merging them into a shapefile, indicating a count of the number of overlaps across all fire layers. We calculated the total area burned by dissolving fire layers for all 20 years into one layer that represented the area burned during that time period; this method represents the total area burned, but does not account for repeated burning of the same area across years. We calculated the percent of total area burned for each STZ by dividing the total STZ area burned by the total STZ area and multiplying by 100. All figures were created using the package ggplot2 (Wickham 2016) within Program R (R Development Core Team 2018). All area calculations and maps were created using ArcMap 10.5 (ESRI 2016).

## RESULTS

Overall, the seven ecoregions that contribute to Cold Deserts within the US varied in total size and area of fire occurrence from 2000–2019: (Area - % burned) Columbia Plateau (83,131 km^2^ - 13%), Wyoming Basin (132,682 km^2^ - 3%), Northern Basin and Range (140,198 km^2^ - 38%), Snake River Plain (53,626 km^2^ - 13%), Central Basin and Range (308,791 km^2^ - 28%), Colorado Plateaus (136,576 km^2^ - 4%), and Arizona/New Mexico Plateau (146,859 km^2^ - 1%). Below, we present results for the PSTZs created by Bower et al. (2014) stratified by ecoregion.

### Question 1 - Wildfire occurrence within Cold Desert ecoregions of the US

#### Frequency of Fire

Most areas that burned within Cold Deserts during our focal 20-year timeframe only burned one time, accounting for 83.5% of total area burned (Fig. 2A). Areas that burned twice during that time frame accounted for 13.5% of total area burned, followed by burning three times (2.3%), four times (0.6%), and five or more times (0.06%). The Northern Basin and Range experienced the most area burned, with 28,898 km^2^ burning one time, 4,350 km^2^ burning two times, 880 km^2^ burning three times, and 383 km^2^ burning four or more times from 2000–2019 (Fig. 3). This was followed closely by the Central Basin and Range, where more area burned a single time (31,007 km^2^), but there was less area experiencing repeated burning (Fig. 3). Only small areas within four ecoregions experienced fire five or more times within the focal time frame: Northern Basin and Range (47 km^2^), Central Basin and Range (17 km^2^), Columbia Plateau (2 km^2^), Snake River Plain (1 km^2^). The three ecoregions where fire was less frequent experienced less repeated burning overall, relative to total area burned: Arizona/New Mexico Plateau (17%), Colorado Plateaus (2%), and Wyoming Basin (2%).

**Figure 2.**
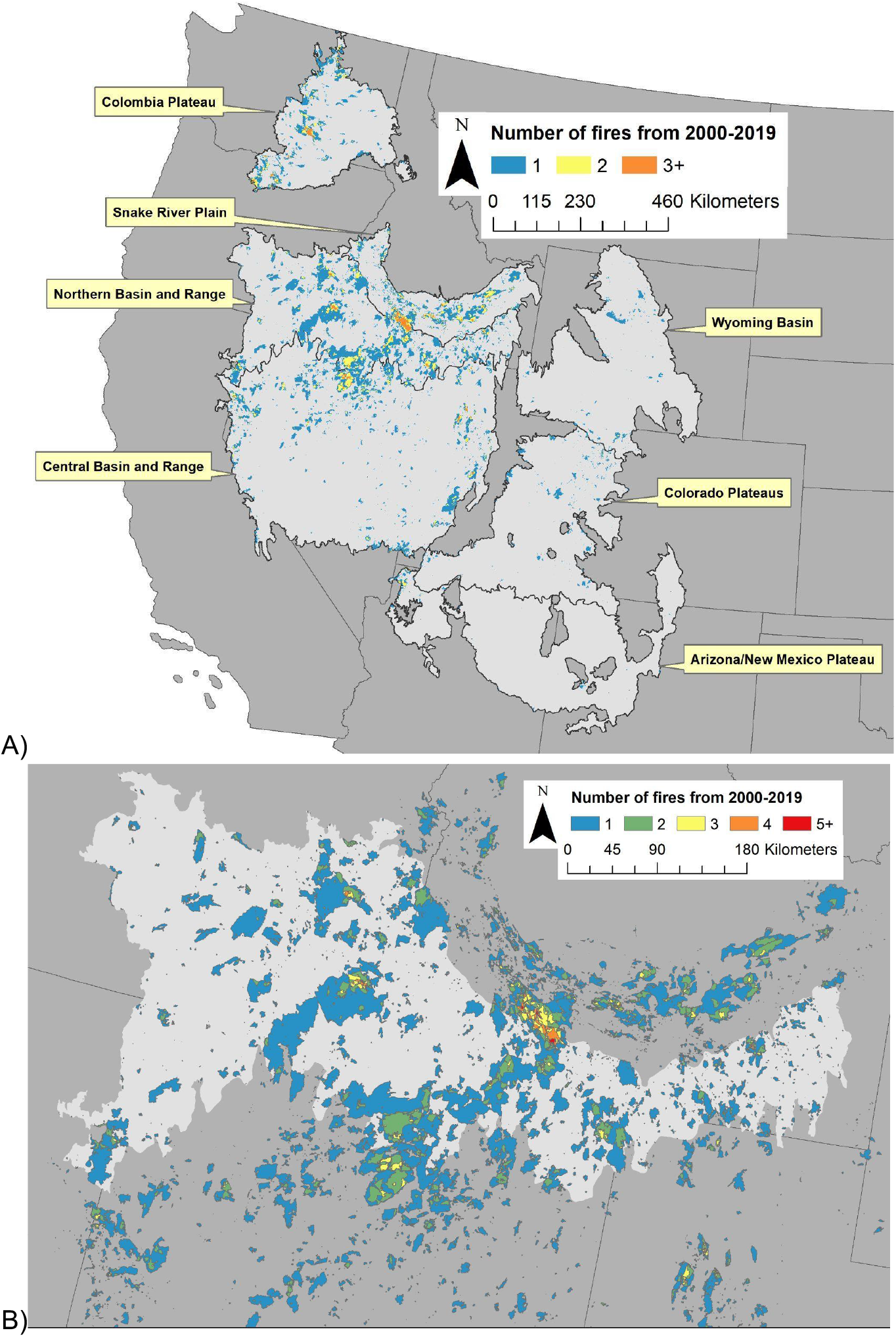
Maps showing burn frequencies of different areas within A) areas of Cold Desert showing the different Omernik Level III ecoregions and B) a close-up of the Northern Basin and Range Ecoregion, with cooler colors representing less frequent burning and hotter colors representing more frequent burning.

**Figure 3.**
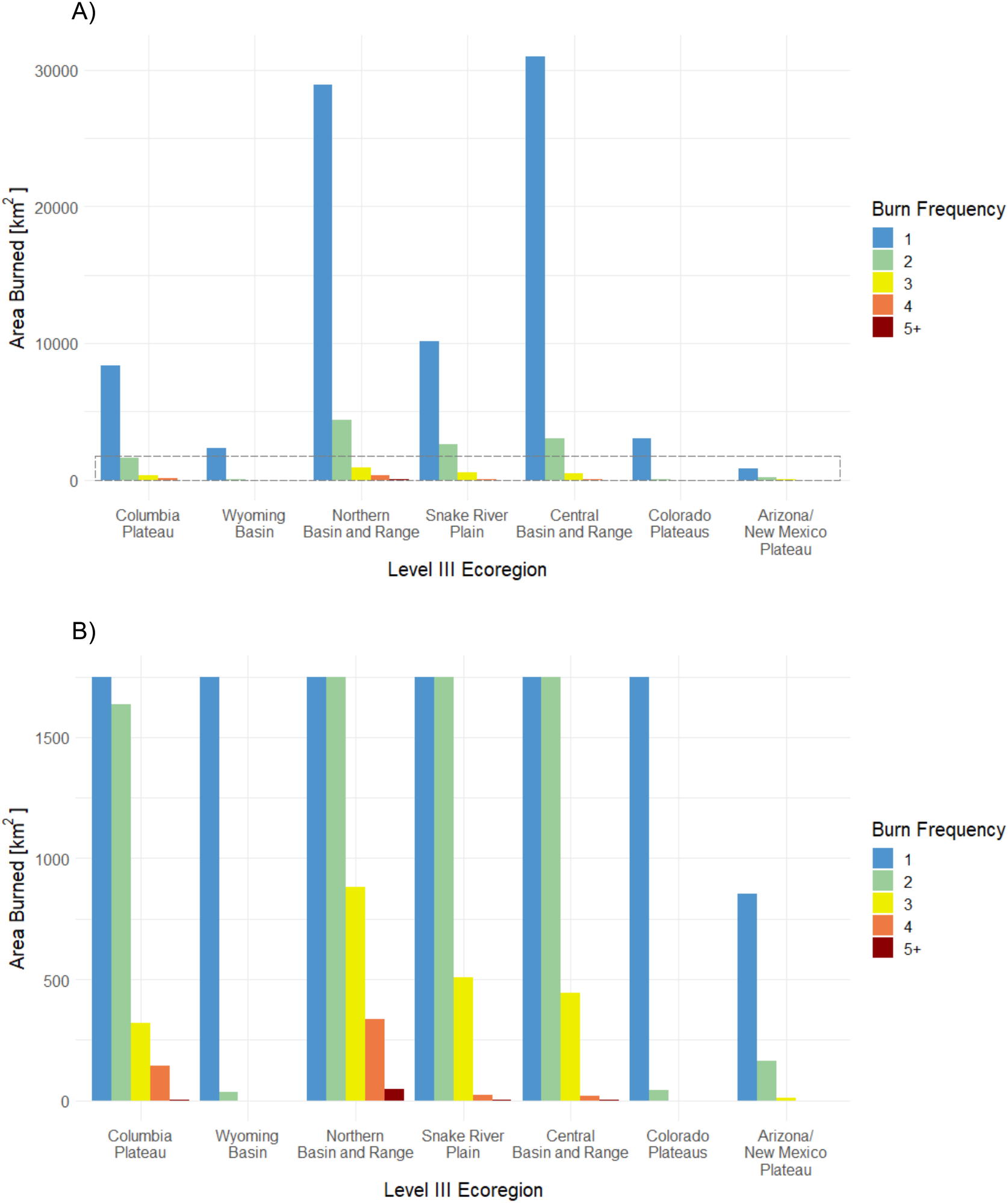
Summary of burn frequencies for level III ecoregions from 2000–2019 showing A) the total area burned for each burn frequency within each ecoregion and B) a zoomed-in view of the area within the dashed box in panel A, showing a y-axis cut-off at 1750 km^2^, to accentuate areas experiencing higher burn frequencies. Note that the total area of each ecoregion differs, see Results.

#### Total Area Burned within Provisional Seed Zones (PSTZs)

Ecoregions varied in which PSTZs were burning the most, as measured by total area burned (Fig. 4, Table 2). In general, areas with an annual heat:moisture index (AH:M) of 6–12 burned the most, with the exception of the Colorado Plateaus ecoregion and the Wyoming Basin ecoregion where areas with an AH:M of 3–6 burned the most (Figs. 4B and 4F). Though there are 132 PSTZs across all ecoregions within the Cold Deserts as a whole, a majority of the burned area in each ecoregion occurred within two PSTZs, ranging from 50.1% in the Snake River Plain to 69.1% in the Columbia Plateau (Table 2). In addition, greater than 82% of the burned area within each ecoregion occurred within five PSTZs (ranging from 82.5% in the Arizona/New Mexico Plateau to 95.8% in the Wyoming Basin (Fig. 4)).

**Table 2.** Provisional seed transfer zones (PSTZs; Bower et al. 2014) with the greatest total area burned for each ecoregion. For each ecoregion, two PSTZs encompass at least 50% of the total area burned. The percent of total area burned within each ecoregion for each listed PSTZ and the number of years (out of a possible 20) the ecoregion burned is in parentheses. Bower et al. (2014) delineated discrete PSTZs for the continental US using a combination of winter minimum temperature and aridity (see Methods for more information).

| Ecoregion | Provisional seed transfer zones most burned |  |  |  | Total % area burned, summed for the two most burned PSTZs |
| --- | --- | --- | --- | --- | --- |
| Columbia Plateau | 20–25F/6–12 | (43% - 19 years) | 25–30F/6–12 | (26% - 20 years) | 69% |
| Wyoming Basin | 10–15F/3–6 | (30% - 20 years) | 10–15F/6–12 | (29% - 20 years) | 59% |
| Northern Basin and Range | 15–20F/6–12 | (45% - 20 years) | 20–25F/6–12 | (19% - 20 years) | 64% |
| Snake River Plain | 20–25F/6–12 | (26% - 20 years) | 15–20F/6–12 | (24% - 20 years) | 50% |
| Central Basin and Range | 15–20F/6–12 | (43% - 20 years) | 20–25F/6–12 | (17% - 20 years) | 60% |
| Colorado Plateaus | 10–15F/3–6 | (38% - 20 years) | 15–20F/3–6 | (23% - 20 years) | 61% |
| Arizona/New Mexico Plateau | 25–30F/6–12 | (31% - 19 years) | 30–35F/6–12 | (23% - 16 years) | 54% |

**Figure 4.**
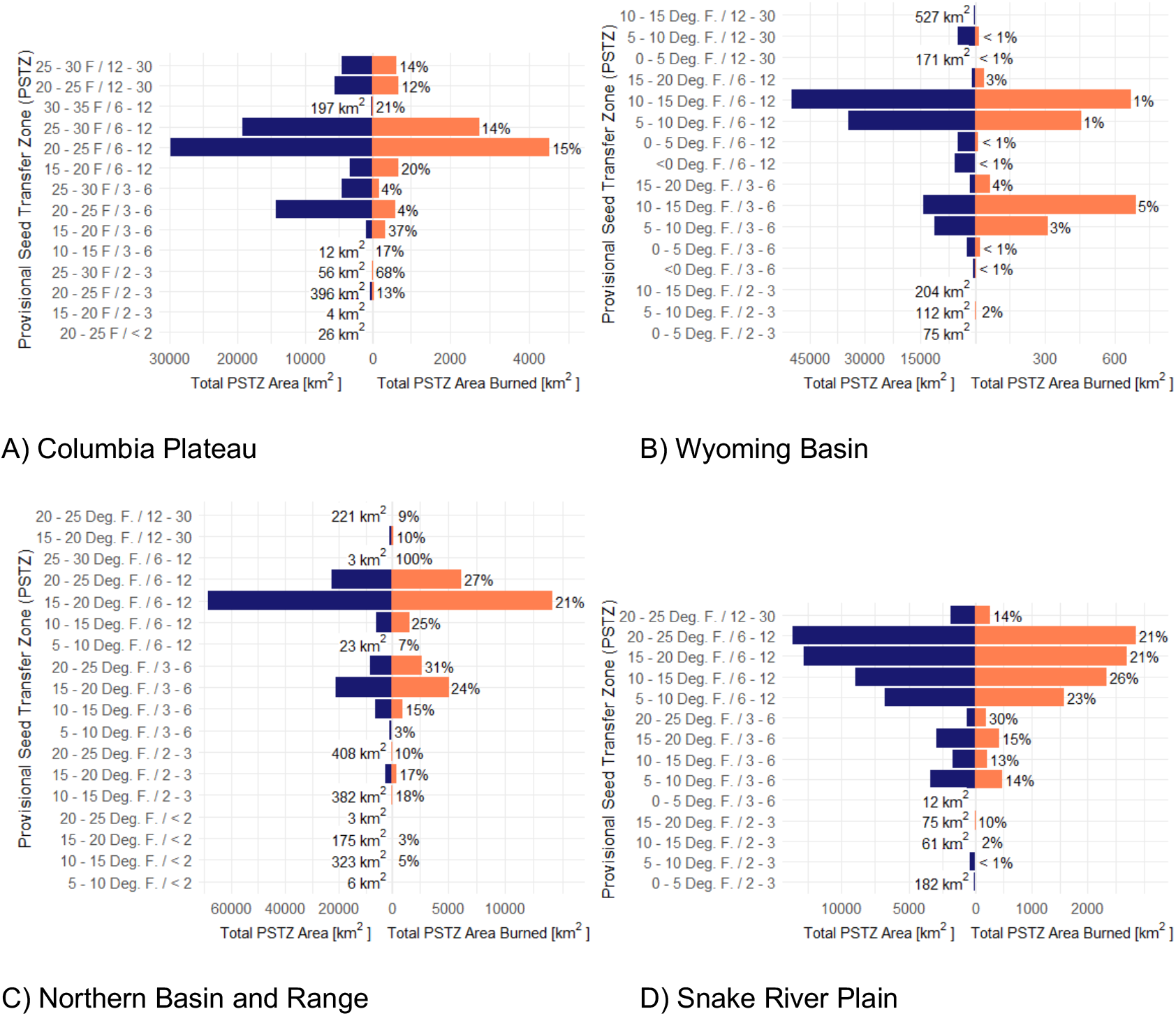
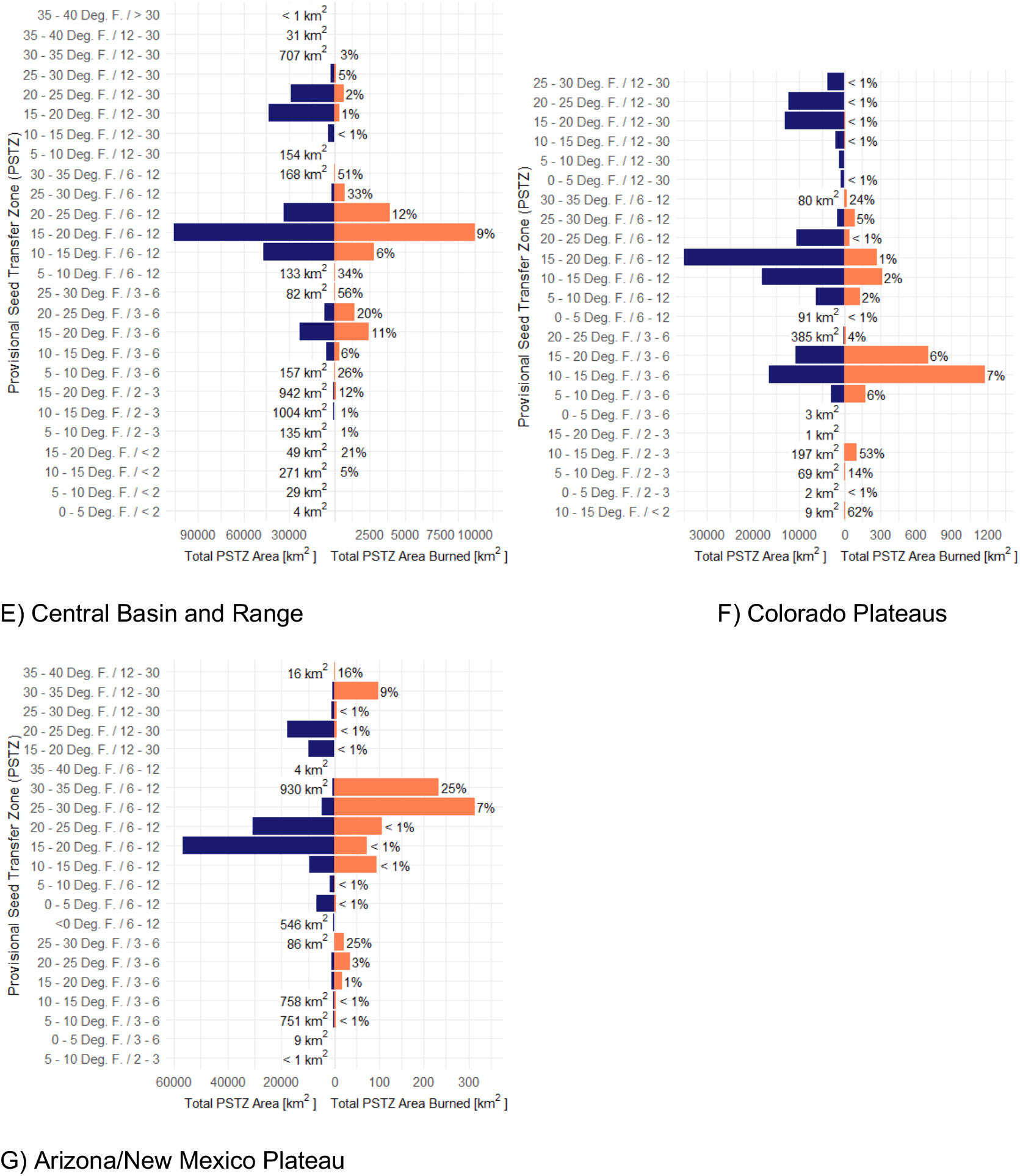
Total area (blue, left) and area burned (orange, right) for each provisional seed transfer zone (PSTZ) within each Cold Desert ecoregion. Notice different x-axis scales for Total PSTZ Area, on the left, and Total PSTZ Area Burned, on the right. The total area of a PSTZ is presented numerically on the left when the value was too small to be visible as a blue bar. Percentages shown to the right of the orange bars indicate the percent of the total PSTZ area burned; when a percentage is missing, that indicates that the area did not burn. Bower et al. (2014) delineated discrete PSTZs for the continental US using a combination of winter minimum temperature and aridity (see Methods for more information).

Sometimes the total area burned was proportional to the size of the ecoregion. For example, in the Northern Basin and Range and the Snake River Plain, the total area burned within different PSTZs was proportionate to their total area (Figs. 4C and 4D). However, in other ecoregions, like the Colorado Plateaus and the Arizona/New Mexico Plateau, there was no consistent relationship between total PSTZ area and the total area burned (Figs. 4F and 4G).

#### Percent of Total PSTZ Area Burned

Ecoregions varied in which PSTZs burned at a high proportion, as measured by total area burned across all years (Fig. 4). There was only one PSTZ where 100% of the area burned in a single fire, Northern Basin and Range 25– 30F/6–12. There were five other PSTZs where greater than 50% of the area burned over the 20-year observation period. For two of these PSTZs, Columbia Plateau 25–30F/2–3 (68%) and Colorado Plateaus 10–15F/<2 (62%), the fires took place in 1 or 2 years, whereas in the others, Central Basin and Range 25–30F/3–6 (56%), Colorado Plateaus 10–15F/2–3 (53%), and Central Basin and Range 30–35F/6–12 (51%), the cumulative burning took place in fires that occurred in 5–7 different years.

Across all 132 PSTZs, 21 PSTZs had no fires within our focal 20-year timeframe. For PSTZs which did experience fire, 29 experienced burning in less than 1% of their area, 36 experienced burning in 1–10% of their area, and 40 experienced burning in 10–50% of their area.

### Question 2 - Ecoregional Case Study - Occurrence of wildfire in the Northern Basin and Range

#### Frequency of Fire

As mentioned above, the Northern Basin and Range was the Cold

Desert ecoregion that experienced the most area burned from 2000–2019. Within the Northern Basin and Range, 83% of the burned area only burned one time, 13% burned twice, and 4% burned three or more times in this 20-year timeframe (Fig. 3B). Two PSTZs contained a majority of the area that burned three or more times, 15–20F/6–12 (51%) and the 20–25F/6–12 (40%). These two PSTZs also contained more than half of the area burning only a single time, 15–20F/6–12 (43%) and 20–25F/6–12 (19%). This makes sense, given that these two PSTZs make up a high proportion of the total area within the Northern Basin and Range ecoregion (Fig. 4C).

#### Total Area Burned within Empirical Seed Zones (ESTZs)

Using ESTZs delineations for both *A. tridentata* and *P. spicata*, we found that one or a few ESTZs made up a majority of the total area burned for these species.

For example, for *A. tridentata*, ESTZ 21 made up 87% of the total *A. tridentata* area burned within the Northern Basin and Range ecoregion, far exceeding the area burned in all other ESTZs within this region (Figs. 5B and 6A). Further, this ESTZ burned every year from 2000–2019, with an average of 1000 km^2^ burning across all fire years and a range from 2 km^2^ in 2009 to 4,117 km^2^ in 2012.

**Figure 5.**
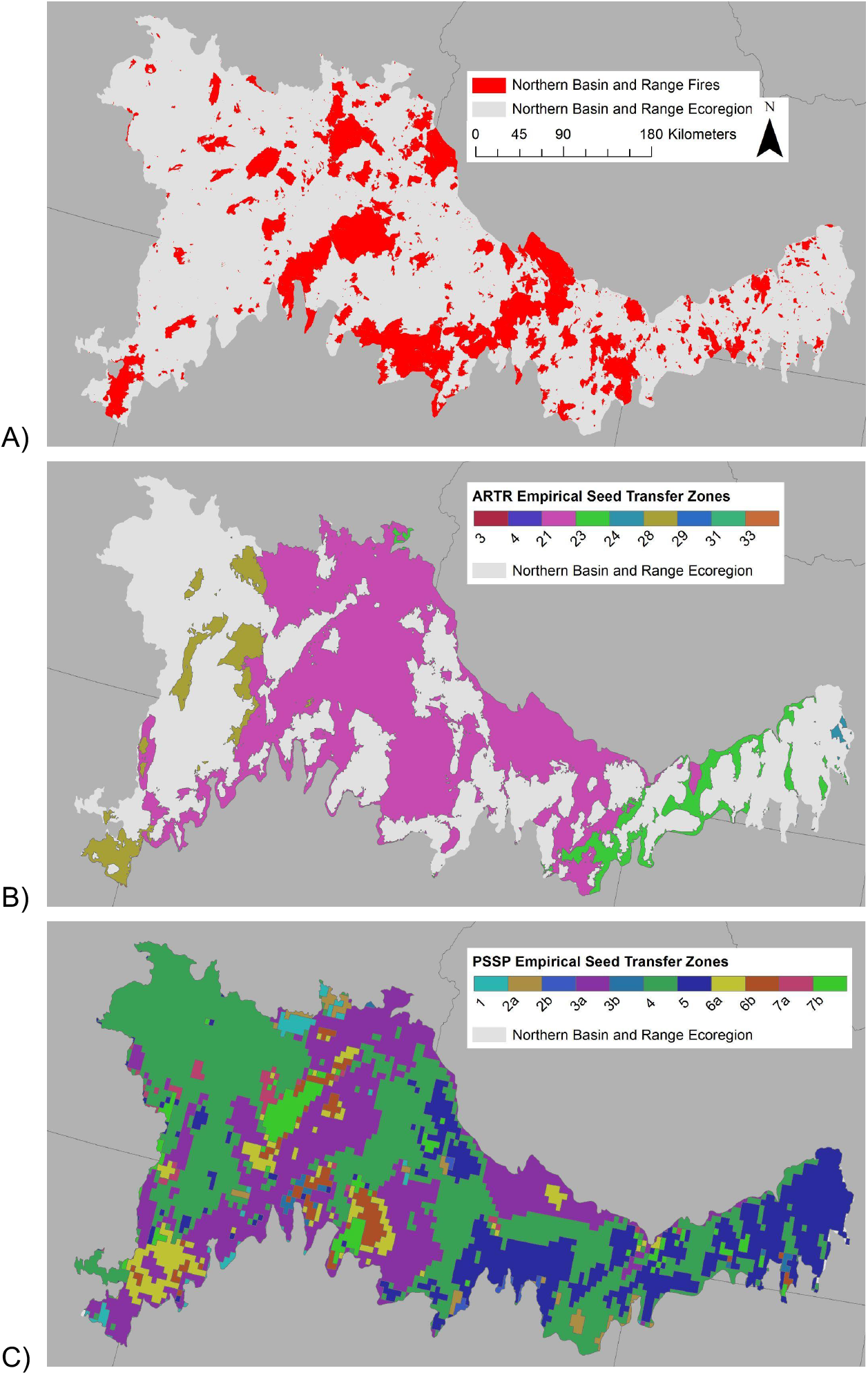
Maps of the Northern Basin and Range ecoregion showing A) areas that have burned from 2000–2019 and empirical seed transfer zones (ESTZs) for B) *Artemisia tridentata* - ssp. *wyomingensis* and *tridentata* (ARTR) and C) *Pseudoroegneria spicata* (PSSP).

**Figure 6.**
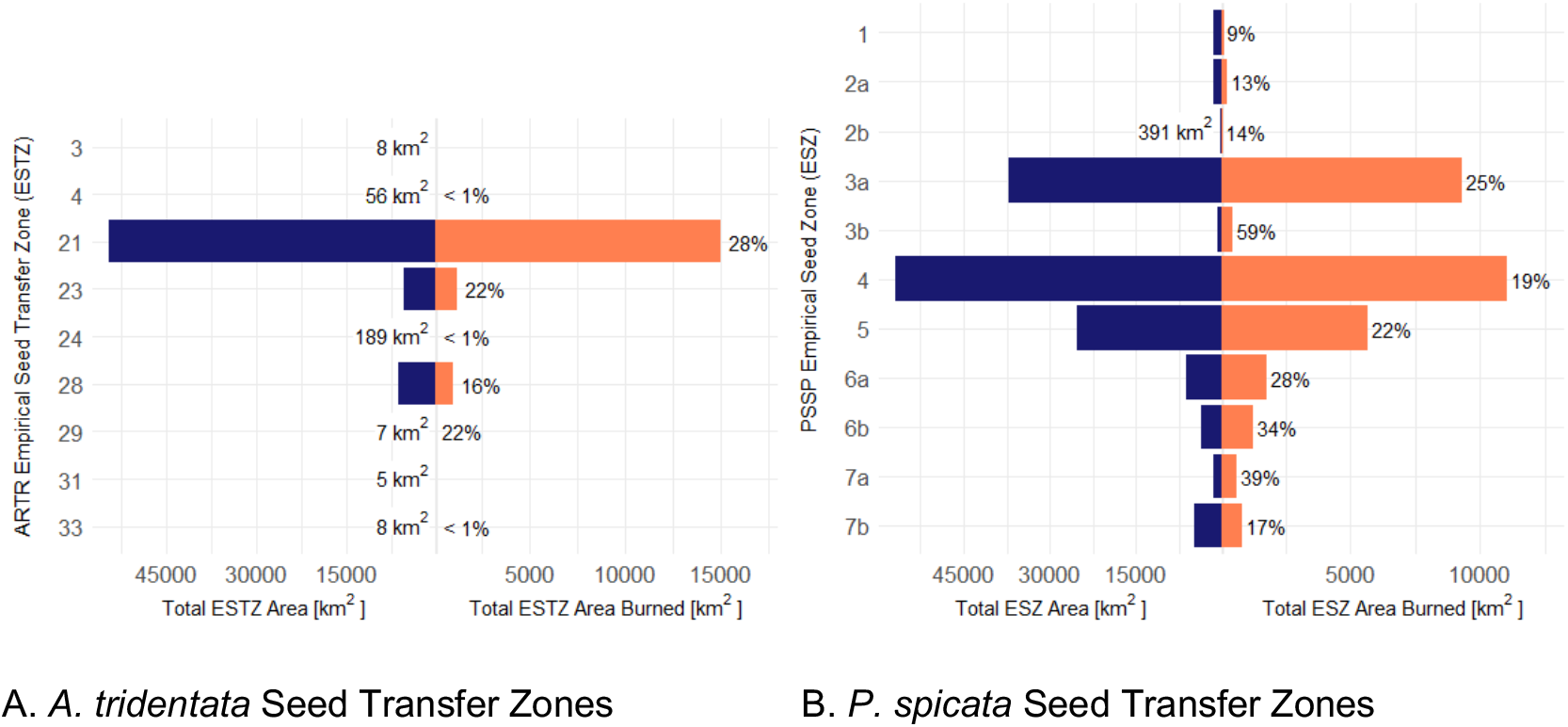
Total area (blue, left) and area burned (orange, right) within the Northern Basin and Range ecoregion for A) empirical seed transfer zones (ESTZ) for sagebrush (*Artemisia tridentate* ssp. *wyomingensis* and *tridentata*) and B) empirical seed transfer zones for bluebunch wheatgrass (*Pseudoroegneria spicata*). Notice different x-axis scales for Total Seed Transfer Zone Area, on the left, and Total Seed Transfer Zone Area Burned, on the right. The total area of an ESTZ is presented numerically on the left when the value was too small to be visible as a blue bar. Percentages shown to the right of the orange bars indicate the percent of the total ESTZ area burned; when a percentage is missing, that indicates that the area did not burn.

Burned areas spanned more ESTZs for *P. spicata*, but some zones had much greater fire activity than others. For example, ESTZ 4 made up 35%, and ESTZ 3a made up 30% of the total *P. spicata* ESTZ area burned within the Northern Basin and Range ecoregion (Figs. 5C and 6B). Both of these ESTZs burned every year from 2000–2019, with an average of 669 km^2^ in ESTZ 4 and an average of 613 km^2^ in ESTZ 3a burning across all fire years. Fires ranged in size from 2–2,846 km^2^ in ESTZ 4 and from 2–2,694 km^2^ in ESTZ 3a, with the smallest area burned occurring in 2009 and the largest area burned occurring in 2012 for both ESTZs.

#### Percent of Total ESTZ Area Burned

Using ESTZs delineations for both *A. tridentata* and *P. spicata*, we found that *A. tridentata* had only a few imperiled ESTZs that experienced a high proportion of burning, whereas *P. spicata* experienced a high proportion of burning in several of its ESTZs.

For *A. tridentata*, ESTZ 21 burned at a high proportion (28%, Fig. 6A), with an average of 2% of the ESTZ area burning across all years and a range from nearly 0% in 2009 to 8% in 2012. Two smaller ESTZs, ESTZ 23 and ESTZ 29, also experienced burning at a relatively high proportion, 22% (Fig. 6A). For ESTZ 23, an average of 1% of its area burned across all years, with a range of 0% in 2004 to 8% in 2007. ESTZ 29 is a relatively small ESTZ (7 km^2^) that also burned in a relatively high proportion; this ESTZ only burned twice, with 1% burning in 2007 and 21% burning in 2016.

Six *P. spicata* ESTZs experienced fire across 25% or more of their area within the 20-year timeframe (Fig. 6B). Two of the ESTZs, ESTZ 3a and ESTZ 5, are relatively large (>25,000 km^2^) and experienced fire every year, with an average of 1–2% of the ESTZ area burning across all 20 years and a range from nearly 0% to 7% of the area burning in any individual year. The remaining four ESTZs are all relatively small (<6,500 km^2^). ESTZ 3b burned nine years out of the 20-year time period, with a majority of the burned area occurring in one year (2012), when 54% of the ESTZ burned. For ESTZ 6a, fire occurred every year during our focal timeframe, with an average of 2% burning across all years and a range from nearly 0% (2008) to 13% (2012). ESTZ 6B showed a similar pattern, with an average of 2% burning across all years and a range from 0% (2008) to 18% (2012). ESTZ 7a burned 13 years out of the 20-year time period, with an average of 3% burning across all years with fire and 25% of the ESTZ area burning in a single year (2012).

### Question 3 - Greater Sage-Grouse Case Study - Occurrence of wildfire in Priority Areas for Conservation (PACs) within the Northern Basin and Range

Overlaying the boundaries of the priority greater sage-grouse habitat areas (Figure S1A) within the Northern Basin and Range ecoregion (Figure S1B) allowed us to ask whether particular STZs could be prioritized for restoration in these areas of overlap, and whether results were appreciably different for PACs than for the ecoregion as a whole. We found that fire frequencies and total area burned for both PSTZs and ESTZs were very similar in prime habitat and the entire ecoregion, as were calculations of the percent of each STZ burned (Figure S2A and S2B). There were some small differences in the proportion of ESTZs burned for *A. tridentata* when considering the PACs or entire ecoregion. For example, 32% of ESTZ 28 burned in the PACs vs. 16% in the ecoregion as a whole (Figure S2C). Similarly, for *P. spicata*, the larger ESTZs experienced fire more frequently (19–20 years) and in comparable proportion to the entire ecoregion, with the exception of ESTZ 5; 32% of this zone burned in the PACs area, relative to 22% in the entire ecoregion (Figure S2D).

## DISCUSSION

Seed transfer zones can make restoration more effective, as they indicate areas where seeding is likely to be successful (Kilkenny 2015). However, seed zones present practical challenges as well, by introducing complexity into the collection, agricultural production, and purchase of native seeds (Nevill et al. 2016). Here, we sought to simplify this process, asking whether we could identify priority seed zones by using geospatial information from historical fire records to identify likely seed needs. Our hope was that historical data could be used to anticipate restoration and procurement needs so that seed is ready to use within the tight timelines imposed by practical and ecological constraints (Erickson & Halford 2020). Indeed, while there were multiple STZs represented in each ecoregion, we found that only two PSTZs made up a majority of the total area burned within each ecoregion. Additionally, some of these PSTZs burned every year (23 PSTZ-ecoregion combinations out of a total of 132 examined here). In some cases, larger STZs experienced more burning, but not always; sometimes area burned was higher than expected based on the total STZ extent. Further, past burning (2000– 2009) was predictive of later burning (2010–2019) within Cold Deserts (Figure 1). This indicates that historic fire data and information on the most widely and frequently burned seed zones may help forecast future seed needs if conditions remain similar, and guide seed collection, storage, and production efforts. We also found that considering high-priority conservation goals (in our case, preservation of sage-grouse core habitat) provided only slightly different recommendations for seed procurement than considering the landscape as a whole. Together, our findings support prioritizing restoration activities in the Cold Deserts of the western United States, including seed collection for gene conservation and seed production, procurement, storage, and planning for post-fire seeding and wildlife habitat improvement projects. Additionally, the methods presented here can be adapted for use in other systems around the world, especially in regions where geospatial layers for climate and ecological disturbance are available (e.g., Tareq et al. 2018).

Disturbance size and frequency is changing in response to human activity around the globe (Turner 2010). Historic observations of fire frequency in the Cold Deserts of North America have clear patterns of shorter fire return intervals driven by annual grass invasions and other land use impacts (D’Antonio & Vitousek 1992; Pilliod, Welty & Arkle 2017). As we saw here, the Northern Basin and Range experienced three or more fires in 1,263 km^2^ across the 20-year timeframe, making up 3.7% of the total area burned within the ecoregion. These small areas of repeated fire may be driving the perception that fire return intervals are less than 5 years across Cold Deserts (e.g., Whisenant 1990).

However, other estimates of fire return intervals are more modest (e.g., Balch et al. 2013), consistent with what we observed in our study: 83.5% of the area burned across all Cold Desert ecoregions burned a single time during our focal 20-year timeframe. This might seem contradictory to our result demonstrating that the same STZs burned consistently (e.g., Table 2); however, different areas within these STZs were burning each time. This was true both across the Cold Desert as a whole and for the areas included in our case studies. Because of the greater likelihood of retaining native species after a single burn event (Winward 1985), some sites with longer fire return intervals and less overall disturbance may be good targets for restoration activities (Chambers et al. 2014; Svejcar et al. 2017). Conversely, we also identified relatively small areas experiencing repeated burning, which may be more vulnerable to invasion (Bradley et al. 2018); these conditions generally require additional considerations when planning and implementing restoration activities, such as herbicide use or other targeted site preparation methods.

We were also interested in identifying STZs that experienced burning across all or most of their extent to prioritize potentially imperiled ecological communities that could be a target for conservation, which is a crucial step for guiding seed procurement and other actions (Castillo-Mandujano & Smith-Ramírez 2022). In our study, STZs varied in the proportion of wildfire occurrence, with some being much more affected by fire than others. The persistence of these imperiled native plant communities may depend on the availability of appropriate seed, and restoration could benefit from identifying areas for seed banking and gene conservation (Povilitis 2001). In particular, our case study of *A. tridentata* ESTZs illustrated that there was a major increase in the proportion burned within the sage-grouse priority habitat area (32% ESTZ 28 burned within the Northern Basin and Range PAC, relative to 16% in the entire ecoregion), which is important information for wildlife habitat restoration. In other regions, identifying seed zones with a high proportion of disturbance from development, floods, or mining may also be helpful when considering imperilment as a criterion for prioritizing seed production (e.g., Silcock & Fensham 2018). We note, however, that over-interpretations of the fraction burned should be avoided for some of the very small STZs represented in this study. Use of specified boundaries (such as when we reduced our study area to the western United States Cold Desert or reduced it further to priority habitat for sage-grouse) will necessarily generate small sections of STZs that primarily exist outside of the boundaries of interest. These sections will represent only a portion of the area of the associated STZ and are thus potentially subject to sample area bias and edge effects. However, in some cases this may be worth considering, despite the potential for bias.

Our case study focusing on greater sage-grouse habitat (PACs) demonstrates how geospatial data can be used to identify seed needs for specific habitat conservation actions. Native plants are the foundation of intact sagebrush steppe plant communities and provide essential habitat that greater sage-grouse depend on for breeding, nesting, and foraging (Pennington et al. 2016). We found that fire occurrence in greater sage-grouse PACs within the Northern Basin and Range ecoregion reflected a similar pattern to fire occurrence within the ecoregion as a whole, with a few exceptions showing slight shifts in STZ priority or for imperiled STZs. This indicates that restoration planning based on ecoregional data will largely satisfy the specific needs for post-fire restoration in sage-grouse PACs within this ecoregion. Sagebrush steppe plant communities are a focus area for the Bipartisan Infrastructure Law (BIL 2021) and Federal directives encourage that native plant materials be used in seeding (e.g., BLM 2008); our findings provide a way to prioritize restoration to ensure that the right seed is available. In other ecosystems, investigating the alignment of overlapping restoration needs (e.g., regional vs. wildlife habitat) may identify ways to balance the acquisition of appropriate seed with limited resources efficiently.

Once priority seed zones have been identified, acquiring appropriate seed resources can still present challenges, with different considerations for species that are wild-collected vs. farmed, and whether the seed zone is imperiled (representing limited and declining opportunities for wild seed collection; Nevill et al. 2016). For example, in our study region, *A. tridentata* seed is primarily collected from wild populations. However, the volume of *A. tridentata* seed required to meet annual restoration demand may exceed the availability of wild-collected seed for some ESTZs, especially in seed zones that experience more frequent fire. These seed zones tend to have both greater restoration needs and represent environments that, because they are on the more arid end of the species’ range, are less conducive to wildland seed production (Shaw et al. 2005), which may signal the need for alternative avenues of seed procurement, such as the development of seed orchards (Zinnen et al. 2021) and greater long-term storage capacity to take advantage of years with higher wildland seed yields (Merritt & Dixon 2011; Havens et al. 2015). For other commonly used species, such as *P. spicata*, seed collection from wild populations is not feasible for the scale of restoration application (Pilliod, Welty & Toevs 2017), and agricultural seed production is always necessary to meet restoration needs (Pedrini et al. 2020). Identifying priority seed zones is an excellent first step, but even then, seed production fields for long-lived perennials often take several years to achieve adequate production levels (Shock et al. 2016). Further, annuals and shorter-lived perennials may also have unique seed production needs and harvesting requirements (de Queiroz et al. 2021). Due to these challenges, production for the commercial market will likely only be viable for seed zones where there is adequate recurring demand, and seed from zones with lower demand will likely only be available through direct contract production (Taylor et al. 2018). Additional seed procurement obstacles may be faced at the global level, especially in developing countries where ecological, resource, and social contexts can pose challenges, including having more biodiverse biomes to manage, growing populations and attendant trade-offs between production of food and other necessities, fewer technological resources for research and decision-making, and dependency on foreign aid to conduct restoration and conservation projects (Alexander et al. 2011; Chazdon and Laestaduis 2016; Fagan et al. 2020). Tools that might forecast seed needs based on historical ecological disturbance patterns, like those presented here, can help restoration practitioners in these regions focus efforts and leverage limited resources to develop proactive native seed markets that support restoration priorities.

While many ecological disturbances are unplanned and somewhat unpredictable, data on past disturbance patterns are available for many regions of the world (Pandey et al. 2019) and could become more widely available with increased access to global environmental monitoring systems (e.g., Sundareshwar et al. 2007). Using the past as a guide for the future, the methods presented here may help generate prioritization strategies for seed production, which are likely to be useful for restoration in response to unplanned disturbance. In our focus on patterns of fire over a 20-year timeframe in western United States Cold Deserts, we demonstrated that fire occurrence, while widespread, tends to be concentrated in a small number of STZs, suggesting that prioritization of the use of the genetic resources represented by those STZs could lower the cost and increase the efficiency and success of restoration projects in this region. Focusing on procurement and storage of seeds from the most burned seed zones appears to be a realistic and feasible effort, as a small number of seed zones represent large areas of need. This is a win-win for seed producers, seed collectors, and seed users, and we have created information that has the potential to better match supply to demand. We note, however, that this approach is only effective if past disturbance is predictive of future disturbance, and as the pace of global change increases, further prioritization methods that incorporate complexity and uncertainty may be needed to manage this challenge.

## ACKNOWLEDGEMENTS

Thanks to Nancy Shaw who consulted on early ideas for this study and provided an early review. The Great Basin Native Plant Project, USDI Bureau of Land Management, USDA Forest Service Rocky Mountain Research Station, USDI Fish and Wildlife Service, and the University of Nevada, Reno provided funding.

The findings and conclusions in this article are those of the authors and do not necessarily represent the views of the U.S. Fish and Wildlife Service; the findings and conclusions in this publication are those of the authors and should not be construed to represent an official USDA or US Government determination or policy.

## DATA AVAILABILITY STATEMENT

Geospatial data for ecoregional boundaries can be found at: https://www.epa.gov/eco-research/ecoregions-north-america

Geospatial data for fire perimeters can be found at:

https://data-nifc.opendata.arcgis.com/search?tags=Category%2Chistoric_wildlandfire_opendata

## LITERATURE CITED

Alexander S, Nelson CR, Aronson J, Lamb D, Cliquet A, Erwin KL, Finlayson CM, de Groot RS, Harris JA, Higgs ES, Hobbs RJ, Lewis RRR, Martinez D, Murcia C (2011) Opportunities and Challenges for Ecological Restoration within REDD+. Restoration Ecology 19: 683–689

Arkle RS, Pilliod DS, Hanser SE, Brooks ML, Chambers JC, Grace JB, Knutson KC, Pyke DA, Welty JL, Wirth TA (2014) Quantifying restoration effectiveness using multiscale habitat models: Implications for sage-grouse in the Great Basin. Ecosphere 5:1–32

Balch JK, Bradley BA, D’Antonio CM, Gómez-Dans J (2013) Introduced annual grass increases regional fire activity across the arid western USA (1980-2009). Global Change Biology 19:173–183

Baughman OW, Agneray AC, Forister ML, Kilkenny FF, Espeland EK, Fiegener R, Horning ME, Johnson RC, Kaye TN, Ott J, St. Clair JB, Leger EA (2019) Strong patterns of intraspecific variation and local adaptation in Great Basin plants revealed through a review of 75 years of experiments. Ecology and Evolution 9:6259–6275

[BIL] Bipartisan Infrastructure Law (2021) Infrastructure Investments and Jobs Act, H.R. 3684, 117th Congress. https://www.congress.gov/bill/117th-congress/house-bill/3684

[BLM] Bureau of Land Management (2007) Burned Area Emergency Stabilization and Rehabilitation Handbook. Handbook H-1742-1. Available at https://www.blm.gov/sites/blm.gov/files/uploads/Media_Library_BLM_Policy_Handbook_h1742-1.pdf [Accessed December 2021]

[BLM] Bureau of Land Management (2008) Integrated vegetation management handbook. Handbook H-1740-2. Available at https://www.blm.gov/sites/blm.gov/files/uploads/Media_Library_BLM_Policy_Handbook_H-1740-2.pdf [Accessed July 2022]

[BLM] Bureau of Land Management Public Land Statistics Reports, 2015-2020.Volumes 200-205, P-108-5 to P-108-10. Retrieved from https://www.blm.gov/about/data/public-landstatistics

Bower AD, St. Clair JB, Erickson V (2014) Generalized provisional seed zones for native plants. Ecological Applications 24:913–919

Bradley BA, Curtis CA, Fusco EJ, Abatzoglou JT, Balch JK, Dadashi S, Tuanmu MN (2018) Cheatgrass (Bromus tectorum) distribution in the intermountain Western United States and its relationship to fire frequency, seasonality, and ignitions. Biological Invasions 20:1493–1506

Breed MF, Harrison PA, Bischoff A, Durruty P, Gellie NJC, Gonzales EK, Havens K, Karmann M, Kilkenny FF, Krauss SL, Lowe AJ, Marques P, Nevill PG, Vitt PL, Bucharova A (2018) Priority actions to improve provenance decision-making. BioScience 68:510–516

Broadhurst LM, Lowe A, Coates DJ, Cunningham SA, McDonald M, Vesk PA, Yates C (2008) Seed supply for broadscale restoration: maximizing evolutionary potential. Evolutionary Applications 1:587–597

Castillo-Mandujano J, Smith-Ramírez C (2022) The need for holistic approach in the identification of priority areas to restore: a review. Restoration Ecology e13637:1–11

Chambers JC, Bradley BA, Brown CS, D’Antonio C, Germino MJ, Grace JB, Hardegree SP, Miller RF, Pyke DA (2014) Resilience to stress and disturbance, and resistance to Bromus tectorum L. invasion in cold desert shrublands of western North America. Ecosystems 17:360–375

Chazdon RL, Laestadius L (2016) Forest and landscape restoration: toward a shared vision and vocabulary. American Journal of Botany 103:1869–1871

St. Clair JB, Kilkenny FF, Johnson RC, Shaw NL, Weaver G (2013) Genetic variation in adaptive traits and seed transfer zones for Pseudoroegneria spicata (bluebunch wheatgrass) in the northwestern United States. Evolutionary Applications 6:933–948

D’Antonio CM, Vitousek PM (1992) Biological invasions by exotic grasses, the grass/fire cycle, and global change. Annual Review of Ecology and Systematics 23:63–87

Dumroese RK, Luna T, Richardson BA, Kilkenny FF, Runyon JB (2015) Conserving and restoring habitat for Greater Sage-Grouse and other sagebrush-obligate wildlife: the crucial link of forbs and sagebrush diversity. Native Plants Journal 16:276–299

Erickson VJ, Halford A (2020) Seed planning, sourcing, and procurement. Restoration Ecology 28:S219–S227

[ESRI] Environmental Systems Research Institute (2016) ArcGIS Desktop: Release 10.5.

Fagan ME, Reid JL, Holland MB, Drew JG, Zahawi RA (2020) How feasible are global forest restoration commitments? Conservation Letters 13: e12700

Fulé PZ (2008) Does it make sense to restore wildland fire in changing climate? Restoration Ecology 16:526–531

Golos PJ, Dixon KW, Erickson TE (2016) Plant recruitment from the soil seed bank depends on topsoil stockpile age, height, and storage history in an arid environment. Restoration Ecology 24:S53–S61

Harrison SP, Schoen R, Atcitty D, Fiegener R, Goodhue R, Havens K, House CC, Johnson RC, Leger E, Lesser V, Opsomer J, Shaw N, Soltis DE, Swinton SM, Toth E, Young SA (2020) Preparing for the need for a supply of native seed. Ecological Restoration 38:203–206

Havens K, Vitt P, Still S, Kramer AT, Fant JB, Schatz K (2015) Seed sourcing for restoration in an era of climate change. Natural Areas Journal 35:122–133

Johnson R, Stritch L, Olwell P, Lambert S, Horning ME, Cronn R (2010) What are the best seed sources for ecosystem restoration on BLM and USFS lands? Native Plants Journal 11:117–131

Kachergis E, Rocca ME, Fernandez-Gimenez ME (2011) Indicators of ecosystem function identify alternate states in the sagebrush steppe. Ecological Applications 21:2781–2792

Kilkenny FF (2015) Genecological approaches to predicting the effects of climate change on plant populations. Natural Areas Journal 35:152–164

Knick ST, Dobkin DS, Rotenberry JT, Schroeder MA, Vander Haegen WM, Van Riper C (2003) Teetering on the edge or too late? Conservation and research issues for avifauna of sagebrush habitats. The Condor 105:611–634

Knutson KC, Pyke DA, Wirth TA, Arkle RS, Pilliod DS, Brooks ML, Chambers JC, Grace JB (2014) Long-term effects of seeding after wildfire on vegetation in Great Basin shrubland ecosystems. Journal of Applied Ecology 51:1414–1424

Krawchuk MA, Moritz MA, Parisien MA, Van Dorn J, Hayhoe K (2009) Global pyrogeography: The current and future distribution of wildfire. PLoS ONE 4:1–12

Larson KB, Tuor AR (2021) Deep Learning Classification of Cheatgrass Invasion in the Western United States Using Biophysical and Remote Sensing Data. Remote Sensing 13: rs13071246

Leger EA, Baughman OW (2015) What seeds to plant in the great basin? Comparing traits prioritized in native plant cultivars and releases with those that promote survival in the field. Natural Areas Journal 35:54–68

Liu Z, Wimberly MC (2016) Direct and indirect effects of climate change on projected future fire regimes in the western United States. Science of the Total Environment 542:65–75

Mahood AL, Balch JK (2019) Repeated fires reduce plant diversity in low-elevation Wyoming big sagebrush ecosystems (1984–2014). Ecosphere 10:1–19

Massatti R, Prendeville HR, Larson S, Richardson BA, Waldron B, Kilkenny FF (2018) Population history provides foundational knowledge for utilizing and developing native plant restoration materials. Evolutionary Applications 11:2025–2039

McArthur ED, Fairbanks DJ (2001) Shrubland ecosystem genetics and biodiversity: proceedings. In: Wildland Shrub Symposium: June 13-15, 2000. USDA Forest Service - Rocky Mountain Research Station, Provo, UT p. 365.

McKay JK, Christian CE, Harrison S, Rice KJ (2005) ‘How local is local?’ - A review of practical and conceptual issues in the genetics of restoration. Restoration Ecology 13:432–440

Merritt DJ, Dixon KW (2011) Restoration Seed Banks - a matter of scale. Science 332:424–425

Nafus AM, Svejcar TJ, Ganskopp DC, Davies KW (2015) Abundances of coplanted native bunchgrasses and crested wheatgrass after 13 years. Rangeland Ecology and Management 68:211–214

[NNSP] Nevada Native Seed Partnership (2020) Nevada Seed Strategy. Accessed July 2022: http://agri.ng.gov/uploadedFiles/agrinvgov/Content/Plant/Seed_Certification/FINALStrategy_with%20memo_4_24_20_small.pdf

Nevill PG, Tomlinson S, Elliott CP, Espeland EK, Dixon KW, Merritt DJ (2016) Seed production areas for the global restoration challenge. Ecology and Evolution 6:7490–7497

Nichols L, Shinneman DJ, McIlroy SK, de Graaff MA (2021) Fire frequency impacts soil properties and processes in sagebrush steppe ecosystems of the Columbia Basin. Applied Soil Ecology 165:1–12

Noss RF, LaRoe ETI, Scott JM (1995) Endangered Ecosystems of the United States: a preliminary assessment of loss and degradation.

Omernik JM (1987) Ecoregions of the Conterminous United States. Annals of the Association of American Geographers 77:118–125

Ott JE, Kilkenny FF, Summers DD, Thompson TW (2019) Long-Term Vegetation Recovery and Invasive Annual Suppression in Native and Introduced Postfire Seeding Treatments. Rangeland Ecology and Management 72:640–653

Ott JE, Kilkenny FF, Summers DD, Thompson TW, Petersen SL (2022) Post-fire succession of seeding treatments in relation to reference communities in the Great Basin. Applied Vegetation Science 25:1–16

Pandey V, Srivastava PK, Petropoulos GP (2019) The Contribution of Earth Observation in Disaster Prediction, Management, and Mitigation: a holistic view. In: Techniques for Disaster Risk Management and Mitigation, Geophysical Monograph 244. pp. 47–62.

Pedrini S, Gibson-Roy P, Trivedi C, Gálvez-Ramírez C, Hardwick K, Shaw N, Frischie S, Laverack G, Dixon K (2020) Collection and production of native seeds for ecological restoration. Restoration Ecology 28:S228–S238

Pennington VE, Schlaepfer DR, Beck JL, Bradford JB, Palmquist KA, Lauenroth WK (2016) Sagebrush, Greater Sage-Grouse, and the Occurrence and Importance of Forbs. Western North American Naturalist 76:298–312

Pilliod DS, Welty JL, Arkle RS (2017) Refining the cheatgrass–fire cycle in the Great Basin: Precipitation timing and fine fuel composition predict wildfire trends. Ecology and Evolution 7:8126–8151

Pilliod DS, Welty JL, Toevs GR (2017) Seventy-Five Years of Vegetation Treatments on Public Rangelands in the Great Basin of North America. Rangelands 39:1–9

Pitman AJ, Narisma GT, McAneney J (2007) The impact of climate change on the risk of forest and grassland fires in Australia. Climatic Change 84:383–401

[PCA] Plant Conservation Alliance (2015) National Seed Strategy for Rehabilitation and Restoration 2015-2020. Available at http://www.blm.gov/seedstrategy [Accessed May 2021]

Povilitis T (2001) A case for conserving imperiled plants by ecological area. In: Southwestern rare and endangered plants: Proceedings of the 3rd Conference. Maschinsi, J & Holter, L, editors. Vol. 23 USDA Forest Service - Rocky Mountain Research Station, Flagstaff, AZ pp. 9–12.

Prasse R, Kunzmann D, Schröder R (2010) Development and practical implementation of minimal requirements for the verification of origin of native seeds of herbaceous plants.

de Queiroz T, Swim S, Turner PL, Leger EA (2021) Creating a Great Basin native annual forb seed increase program: lessons learned. Native Plants Journal 22:90–102

R Development Core Team (2018) R: A language and environment for statistical computing. Available at http://www.r-project.org

Richardson BA, Chaney L (2018) Climate-based seed transfer of a widespread shrub: population shifts, restoration strategies, and the trailing edge. Ecological Applications 28:2165–2174

Shaw NL, Lambert SM, Debolt AM, Pellant M (2005) Increasing Native Forb Seed Supplies for the Great Basin. In: National Proceedings: Forest and Conservation Nursery Association 2004. Dumroese, R, Riley, L, & Landis, T, editors. USDA Forest Service - Rocky Mountain Research Station, Medford, OR.

Shock CC, Feibert EBG, Rivera A, Saunders LD, Shaw N, Kilkenny FF (2016) Irrigation requirements for seed production of five Lomatium species in a semiarid environment. HortScience 51:1270–1277

Silcock JL, Fensham RJ (2018) Using evidence of decline and extinction risk to identify priority regions, habitats and threats for plant conservation in Australia. Australian Journal of Botany 66:541–555

Sundareshwar P V, Murtugudde R, Srinivasan G, Singh S, Ramesh KJ, Ramesh R, Michener WK, Mitra a P, Morris JT, Myneni RR, Naja M, Nemani R, Purvaja R, Raha S, Vanan SKS, Sharma M, Subramaniam A, Sukumar R, Twilley RR, Zimmerman PR (2007) Environmental Monitoring Network for India. Science 316:204–205

Svejcar T, Boyd C, Davies K, Hamerlynck E, Svejcar L (2017) Challenges and limitations to native species restoration in the Great Basin, USA. Plant Ecology 218:81–94

Tareq SM, Tauhid Ur Rahman M, Zahedul Islam AZM, Baddruzzaman ABM, Ashraf Ali M (2018) Evaluation of climate-induced waterlogging hazards in the south-west coast of Bangladesh using Geoinformatics. Environmental Monitoring and Assessment 190:1–14

Taylor MH, Bartholet RD, Banerjee S (2018) Recommendations for increasing the supply and lowering the cost of native plant materials in Nevada through strategic support of the native plant materials industry.

Turner MG (2010) Disturbance and landscape dynamics in a changing world. Ecology 91:2833–2849

[USFWS] US Fish and Wildlife Service (2013) Greater Sage-grouse (Centrocercus urophasianus) Conservation Objectives: Final Report. Denver, CO. Available at http://www.sagegrouseinitiative.com/wp-content/uploads/2013/07/USFWS_ConservationObjectives-report.pdf [Accessed July 2022]

Vallentine JF (1989) Range Development and Improvements. 3rd ed. Academic Press Inc., San Diego, CA

Whisenant SG (1990) Changing fire frequencies on Idaho’s Snake River Plains: ecological and management implications. In: Symposium on Cheatgrass Invasion, Shrub Die-off, and Other Aspects of Shrub Biology and Management (1989). USDA Forest Service - Intermountain Research Station, Las Vegas, NV pp. 4–10.

Wickham H (2016) ggplot2: Elegant graphics for data analysis. Springer-Verlag New York, NY. ISBN 978-3-319-24277-4. Available at: http://ggplot2.tidyverse.org

Williams JR, Morris LR, Gunnell KL, Johanson JK, Monaco TA (2017) Variation in Sagebrush Communities Historically Seeded with Crested Wheatgrass in the Eastern Great Basin. Rangeland Ecology and Management 70:683–690

Winward A (1985) Fire in the sagebrush grass ecosystem - the ecological setting. In: Rangeland Fire Effects: a symposium. Sanders, K & Durham, J, editors. Boise, ID pp. 2–6.

Zinnen J, Broadhurst LM, Gibson-Roy P, Jones TA, Matthews JW (2021) Seed production areas are crucial to conservation outcomes: benefits and risks of an emerging restoration tool. Biodiversity and Conservation 30:1233–1256

