## Supplemental material for "Not all seed transfer zones are created equal: Using fire history to identify seed needs in the Cold Deserts of the Western US"

*Question 3 - Greater Sage-Grouse Case Study - Occurrence of wildfire in Priority Areas for Conservation (PACs) within the Northern Basin and Range*

Overlaying the boundaries of the priority greater sage-grouse habitat areas (Fig. S1A) within the Northern Basin and Range ecoregion (Fig. S1B) allowed us to ask whether particular STZs could be prioritized for restoration in these areas of overlap.

Frequency of Fire - Fire frequencies within the Northern Basin and Range PACs area were comparable to the fire frequencies across the Cold Desert area as a whole, with 86% of the PACs area burned only experiencing fire one time and 13% of PACs area experiencing fire two times within our 20-year timeframe (Figs. S1B and S2A).

Total Area Burned within PSTZs and ESTZs - For PSTZs within the PACs area, the area burned within the Northern Basin and Range PACs was roughly proportional to the total area they occupied, with two PSTZs, 15–20F/6–12 and 15–20F/3–6, making up a majority of the total area burned, 43% and 18% respectively (Fig. S2B). This is similar to the results for the PSTZs for the entire ecoregion (Fig. 4C).

The STZs with the greatest burned area were also extremely similar for the two focal species with ESTZs. Specifically, for *A. tridentata*, ESTZ 21 was the most burned within the Northern Basin and Range PACs area, making up 85% of the total ESTZ area burned (Fig. S2C), which is essentially identical to the results for the ESTZ for the entire region (87%, Fig. 6A). Similarly, for *P. spicata*, two ESTZs made up a majority of the total PACs area burned, with ESTZ 4 making up 35% and ESTZ 3a making up 25% of the total ESTZ area burned (Fig. S2D); this was also similar to the total ESTZ area burned for the entire Northern Basin and range (35% - ESTZ 4, 30% - ESTZ 3a).

Percent of Total Seed Transfer Zone (STZ) Area Burned - Finally, to consider which STZ areas are most imperiled, we calculated the percent of each STZ that burned. For PSTZs burning within the Northern Basin and Range PACs, the PSTZs burning in 25% or more of their area from 2000–2019 fell into two groups, PSTZs with larger total area (1,000–13,600 $km^2$) and PSTZs with smaller total area (3–217 $km^2$) (Fig. S2B).

Just like the total area burned calculations, imperiled STZs were similar when we considered either the entire Northern Basin and Range or the PACs area specifically. For example, in the ecoregion as a whole, three of the larger PSTZs experienced fire in all 20 years, 20–25F/3–6, 20–25F/6–12, 15–20F/3–6, and this same pattern was observed when we considered only the PACs areas. These are also the largest of the three PSTZs experiencing a high proportion of fire. These three PSTZs experienced an average of 2% burning across all years, but varied in the range of the proportion of burning from nearly 0% to 21%, 18%, and 7%, respectively. The remaining three larger PSTZs experienced fire less frequently, and there were more differences from the entire Northern Basin and Range: 15–20F/2–3, 11 years in PACs (16 years in the entire Northern Basin and Range); 10–15F/3–6, 16 years (relative to 20 years); 10–15F/6–12, 17 years (relative to 19 years); they experienced an average of 2–3% burning across all fire years with a range from nearly 0% to 11–14% of the PSTZ burning in a given year.

For the smaller PSTZs, two experienced fire in only one year, 25–30F/6–12 and 15–20F/< 2. The other three smaller PSTZs experienced fire in 3–6 years during our focal timeframe, with a majority of burning happening in one or two fire years: 5–10F/3–6 (28% - 2018), 10–15F/2–3 (21% - 2007), 15–20F/12–30 (29% - 2012, 10% - 2017).

The proportion of *A. tridentata* ESTZs burning within PACs was somewhat different than in the entire ecoregion. For example, for *A. tridentata* ESTZs within the Northern Basin and Range PACs, three of the four ESTZs burned in 25% or more of their area from 2000–2019, ESTZ 21

(29%), ESTZ 23 (27%), and ESTZ 28 (32%; Fig. S2C). There were some similarities and some differences compared to the ecoregion as a whole: ESTZ 21 (28%), ESTZ 23 (22%), and ESTZ 28 (16%). These three ESTZs all experienced an average of 2% burning across all fire years, but varied in the range of the proportion of burning from nearly 0% to 7% - ESTZ 21, 11% - ESTZ 23, and 28% - ESTZ 28 in any particular fire year. The ESTZ 21 burned in all 20 years, but ESTZ 23 and ESTZ 28 only burned in 15 out of 20 years.

For *P. spicata* ESTZs within the Northern Basin and Range PACs, six of the ESTZs experienced burning in more than 25% of their area from 2000–2019 (Fig. S2D). Three of the ESTZs (3a, 5, 6a) were relatively large (> 5,000 $km^2$), and three of the ESTZs (3b, 6b, 7a) were relatively small (474–2,535 $km^2$). The larger ESTZs experienced fire more frequently (19–20 years) and in comparable proportion to the entire ecoregion, with the exception of ESTZ 5; 32% of this zone burned in the PACs area, relative to 22% in the entire ecoregion. The larger ESTZs experienced an average of 2% burning across all fire years, with a range from 0% or nearly 0% to 9% - ESTZ 3a, 11% - ESTZ 5, and 14% - ESTZ 6a. The smaller ESTZs experienced fire less frequently, in only 6–14 years. For these smaller zones, the proportion of ESTZ area burned in the PACs areas differed substantially from the entire ecoregion (e.g., 82% of ESTZ 3b burned in the PACs vs. 59% in the entire Northern Basin and Range), but these areas were, overall, quite small (Fig. 6). For the smaller ESTZs, they all experienced a majority of their burning in a single year; 79% of ESTZ 3b, 25% of ESTZ 6b, and 36% of ESTZ 7a all burned in 2012.

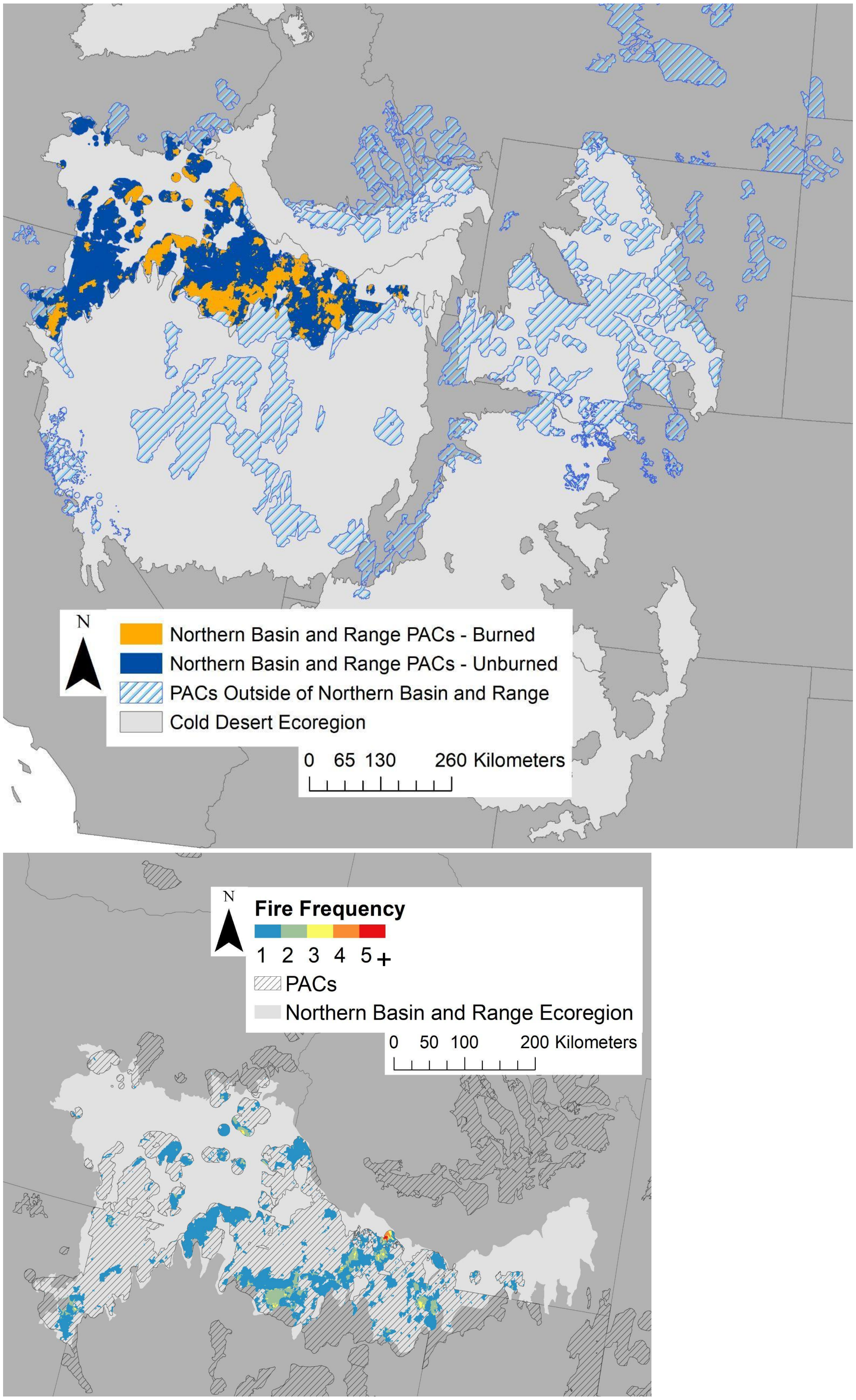


Figure S1. Maps showing A) the Priority Areas of Conservation (PACs) for greater sage-grouse within the western US, and B) the frequency of fire within the Northern Basin and Range PACs.

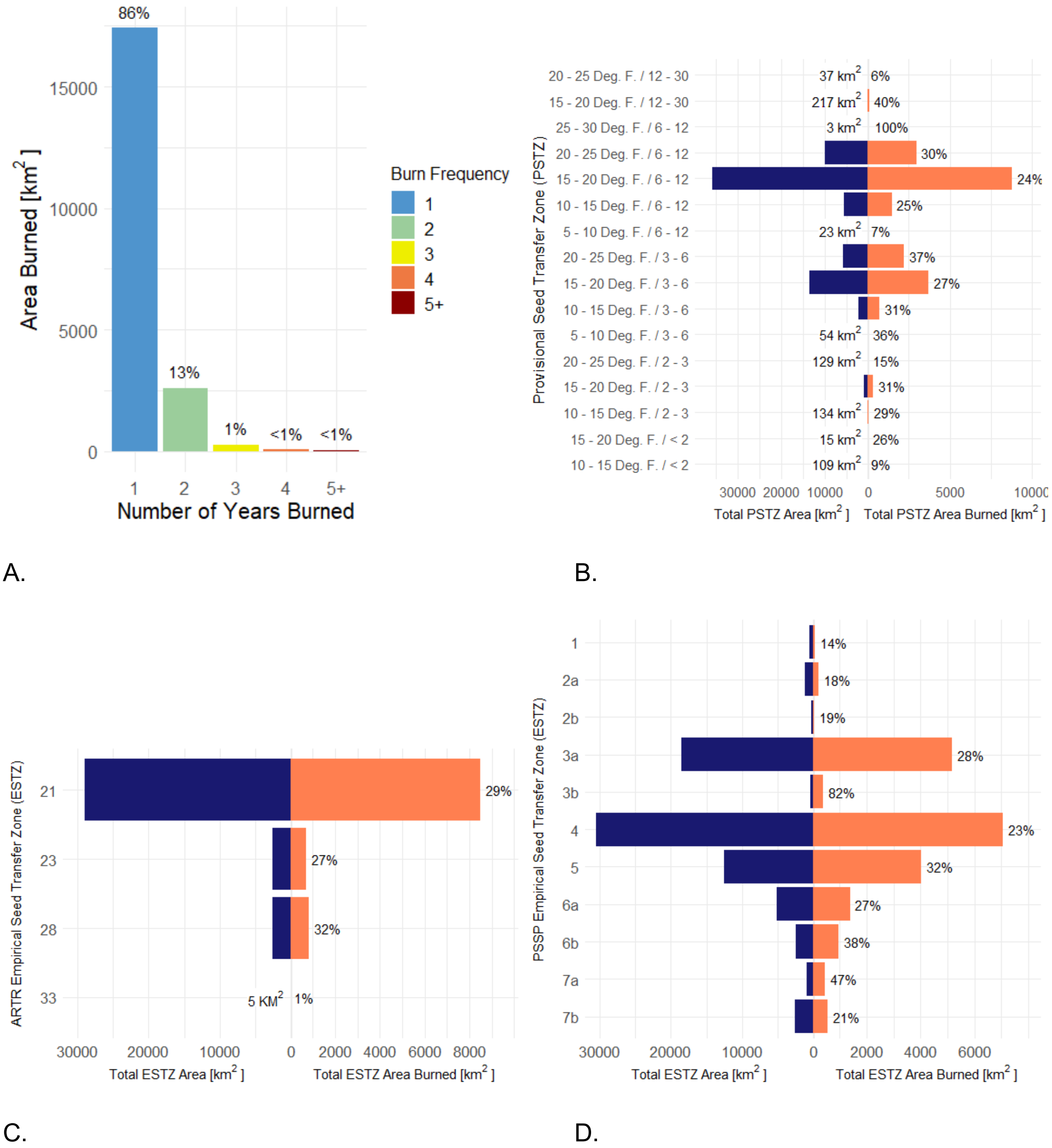


Figure S2. Total area (blue, left) and area burned (orange, right) for Priority Areas of Conservation (PACs) for greater sage-grouse within the Northern Basin and Range ecoregion for A) a summary of the areas experiencing different burn frequencies, B) provisional seed transfer zones (PSTZ), C) empirical seed transfer zones (ESTZ) for sagebrush (*Artemisia tridentata* ssp WY and MT), and D) ESTZs for bluebunch wheatgrass (*Pseudoroegneria spicata*). Notice different x-axis scales for Total Seed Transfer Zone Area, on the left, and Total Seed Transfer Zone Area Burned, on the right. The total area of a particular seed transfer zone is presented numerically on the left when the value was too small to be visible as a blue bar. Percentages shown to the right of the orange bars indicate the percent of the total seed transfer zone area burned; when a percentage is missing, that indicates that the area did not burn. Bower et al. (2014) delineated discrete PSTZs for the continental US using a combination of winter minimum temperature and aridity (see Methods for more information). ESTZs represented here are based on Richardson and Chaney (2018) and St. Clair et al. (2013).
